# Comparative multi-tissue profiling reveals extensive tissue-specificity in transcriptome reprogramming upon cold exposure

**DOI:** 10.1101/2021.03.26.437139

**Authors:** Noushin Hadadi, Martina Spiljar, Karin Steinbach, Gabriela Salinas, Doron Merkler, Mirko Trajkovski

**Author notes:** Equal contribution (Noushin Hadadi, Martina Spiljar).

## Abstract

Cold exposure is an extensively used intervention for enhancing thermogenic and mitochondrial activity in adipose tissues. As such, it has been suggested as a potential lifestyle intervention for body weight maintenance. The metabolic consequences of cold acclimation are not limited to the adipose tissues, however the impact on rest of the tissues in context of their gene expression profile remains unclear. Here we provide a systematic characterization of cold exposure-mediated effects in a comparative multi-tissue RNA sequencing approach using wide range of organs including spleen, bone marrow, spinal cord, brain, hypothalamus, ileum, liver, subcutaneous-, visceral- and brown adipose tissues. Our findings highlight that transcriptional responses to cold exposure exhibit high degree of tissue-specificity both at the gene level and at GO enrichment gene sets, which is not directed by the basal gene expression pattern exhibited by the various organs. Our study places the cold adaptation of individual tissues in a whole-organism framework and provides an integrative transcriptional analysis necessary for understanding the cold exposure-mediated biological reprograming.

## INTRODUCTION

Over the last decade and the discovery of active brown adipose tissue (BAT) presence in humans, cold acclimation for stimulating thermogenesis has gained interest as an intervention leading to increased energy expenditure (Chondronikola et al., 2016; Nedergaard et al., 2007). As such, the association between activating the thermogenic program of BAT and metabolic health has been extensively studied (Walden et al., 2012). Most of the cold-exposure studies are conducted on mouse models, and various interventions differing in length and intensity of the cold have been used to alter adipose tissue activity and metabolism (Peres Valgas da Silva et al., 2019). The BAT is present at distinct anatomical sites, including the interscapular (iBAT), perineal and axillary depots. The white adipose tissue (WAT) stores energy in form of triglycerides and it is found throughout the body. Its largest compartments are the subcutaneous and visceral adipose tissues (SAT and VAT, respectively). Following prolonged cold, brown fat-like cells also emerge in SAT (known as “beige” cells) in a process referred to as fat browning (Chouchani et al., 2019; Stojanovic et al., 2018). Brown and beige fat-associated thermogenesis account for 2-17% of the total 12-20% of energy that is expanded daily (Tan et al., 2011), indicating that other tissues may also contribute (van Marken Lichtenbelt and Schrauwen, 2011), as has recently been shown for the liver (Abumrad, 2017; Simcox et al., 2017). However, the contribution of other organs in the overall adaptation of the organism to cold exposure is less understood (Omran and Christian, 2020).

Hypothalamus is the central regulating unit in the brain for maintenance of the energy homeostasis, including the body temperature. During cold, sympathetic signals are sensed by adipocytes via their beta-adrenergic receptors, which results in BAT activation (Cannon and Nedergaard, 2004; Chechi et al., 2013; Stojanovic et al., 2018). Immune cells are also implicated in the effects of cold exposure in the adipose tissues (Hotamisligil, 2017; Kohlgruber et al., 2016; Molofsky et al., 2013; Omran and Christian, 2020; Qiu et al., 2014; Stojanovic et al., 2018), which harbor an anti-inflammatory immune profile under cold. In obesity, the fat predominantly contains inflammatory immune cells linked to systemic low-grade inflammation (Hotamisligil, 2017; Kohlgruber et al., 2016). The systemic immune state during cold exposure, however, remains largely elusive. While immune cells reside in a variety of organs, the major immune hubs are the primary lymphoid tissues where immune cells are generated (e.g. bone marrow), and the secondary lymphoid tissues where they are activated and expanded (e.g. spleen). Small stretches of lymphoid tissues also reside within other organs, including the intestine that contains the gut-associated lymphoid tissues (GALT). We (Chevalier et al., 2015; Spiljar et al., 2020) and others (Simcox et al., 2017) observed that cold exposure also affects organs apart from the adipose tissues, including the intestine and immunologic tissues. These data suggest that cold exposure exerts a whole-body functional reprogramming, however, no systematic transcriptomics analysis has addressed to what extent organs undergo changes induced by cold exposure. It is also not clear whether various organs display conserved, or tissue specific transcriptional signatures.

RNA sequencing (RNASeq) is the most widely used quantitative approach to assess the global gene expression and its alternations under different conditions, since it determines subtle molecular changes that may contribute to acquisition of certain phenotypes (Carninci et al., 2005; Grada and Weinbrecht, 2013). It has been shown that RNAseq can reflect tissue specificity; as such, highly regulated genes might represent the characteristic functions of tissues (Breschi et al., 2016; Sonawane et al., 2017). However, due to a deluge of data, particularly when several transcriptomic data sets are obtained, the combined analysis of several datasets (meta-analysis) remains challenging (Sudmant et al., 2015). This analysis becomes even more challenging if one aims to interconnect the variations in the meta-data with the complex molecular basis of phenotypic changes. In this study, we conducted a systematic analysis of cold-induced gene expression across ten mouse tissues. We establish a common expression signature of deregulated genes in BAT during various cold exposure experimental setups across seven studies (six previously published RNAseq datasets and this work). Further, we provide a comprehensive resource dataset of comparative transcriptomics across ten mouse tissues describing the transcriptional landscape at room temperature (RT), and its alternations (differential expression) upon cold exposure (CE) (10°C). We systematically investigate how differential expression impacts specific cellular functions across the ten tissues and identify shared, specific, and inversely regulated gene signatures and gene set enrichments under cold. Our work shows that adipose tissues undergo the most pronounced transcriptome deregulations, followed by the immune tissue and central nervous system (CNS) cluster. With this resource and the applied bioinformatics methods, we characterize tissue-specific expression patterns and detect temperature-dependent gene expression profiles; provide insights into the tissue-specific adaptation mechanisms associated with cold exposure; and place the adaptive role of each tissue in a whole-organism perspective to comprehend the tissue-specific organization of the biological processes upon cold.

## RESULTS

### Effectiveness of 10°C for activation and recruitment of the thermogenic capacity of brown adipose tissue (BAT)

We first investigated the effectiveness of a milder cold intervention using 10°C on the BAT following a gradual decrease of the temperature, as an approach that mimics the gradual decrease of the environmental temperature that typically happens in nature. We combined the BAT thermogenic biomarkers introduced in literature (Perdikari et al., 2018) with those genes which are annotated in Gene Ontology (GO) database to any GO term related to adaptive thermogenesis. This resulted in a list of 148 potential thermogenesis-induced marker genes, of which 120 genes were significantly deregulated in at least one of the seven studies (Table S1). We next compared the regulation of these genes in our study to the six other publicly available transcriptomic datasets, in which C57BL/6J male mice were used in a cold exposure intervention (Bai et al., 2017; Cheng et al., 2018; Hao et al., 2015; Marcher et al., 2015; Shore et al., 2013; Xu et al., 2019) (Figure 1A). Several parameters such as gender, initial temperature, the cold temperature, the duration of the cold exposure, and the nature of the exposure (acute or chronic) are varying across these studies (Figure 1A).

**Figure 1:**
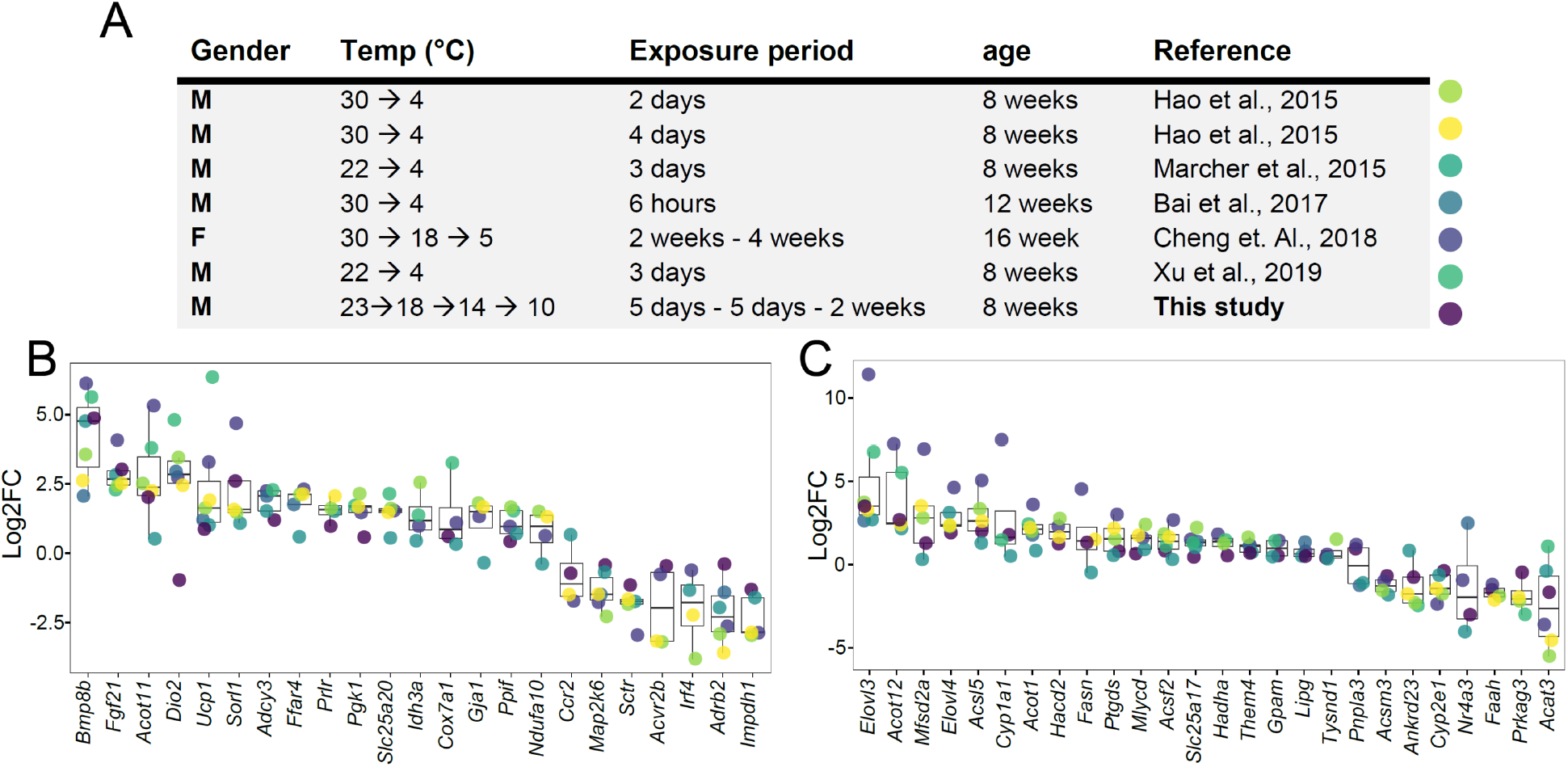
Log2FC of brown adipose tissue (BAT) biomarkers across seven datasets. (**A**) Table of seven cold exposure studies with their corresponding experimental conditions. (**B**) Log2FC of 23 selected BAT thermogenic markers. (**C**) Log2FC of 26 selected fatty acid metabolism gene markers. The markers in (B) and (C) were selected as being significantly (*P*<0.05) regulated in at least four datasets. Each dot corresponds to the log2FC of the indicated gene and colors specify the corresponding study as designated in (A).

Within the identified genes across the seven different studies, we found *Ucp1* and *Bmp8b* being increased in all seven datasets, followed by *Dio2* and *Fgf21* that were regulated in six studies (Table S1). The cold exposure study of Cheng et al. (2018) covered a maximum of 75 of the 120 regulated genes among all the seven studies, follwed by Marcher et al. (2015) with 63 regulated genes and this study with 55 regulated genes, while the study by Bai et al. (2017) identified the least with only 13 dergulated genes. Out of the 120 genes, we selected 23 genes that were significantly regulated in at least four datasets and compared their log2FC across the seven studies (Figure 1B). Despite the notable differences in the experimental conditions, both the number of regulated thermogenesis-induced genes and the magnitude of their regulation in BAT at 10°C were in agreement with those obtained at colder temperatures.

We further focused on 292 genes involved in fatty acid metabolism (as annotated in GO database), a process of critical importance for the cold-induced thermogenesis (Table S2). We identified 190 genes which were significantly deregulated in at least one of the seven studies. Only *Elovl3* was significantly regulated in all seven studies, and we observed a similar trend for the number of fatty acid metabolism-related genes across seven studies, with the maximum number of 127 regulated genes in Cheng et al. (2018), 105 regulated genes in this study and 101 regulated genes in the study by Marcher et al. (2015). The least number of regulated genes was reported in Bai et al. (2017) with 10 regulated genes. Comparing the log2FC of the 26 genes that were significantly regulated in at least four datasets (Figure 1C) shows that consistent with the thermogenesis-regulated genes (Figure 1B), the regulation of fatty acid metabolism related-genes in our study using 10°C is in agreement with the other studies.

These comparisons (both the number of regulated genes and the magnitude of their regulation) suggest that 10°C used in this study induces gene expression alterations that are similar to the harsher cold exposures, indicating that the milder cold temperature is an applicable intervention that resembles the more drastic cold exposures.

### A comprehensive mouse transcriptomic resource across ten tissues at cold exposure

To identify how cold exposure affects the tissue-specific signatures and it whether induces common gene expression changes across various tissues, we performed RNAseq on iBAT, bone marrow, brain, hypothalamus, ileum, liver, inguinal SAT (ingSAT), spinal cord, spleen, and epididymal VAT (epiVAT) from mice exposed to 10°C for two weeks after an acclimatization at 18°C and 14°C for 5 days each and compared them to room temperature-kept controls (Figure 2A). All samples were subjected to quality control and clustering analysis, and we restricted our analyses to all genes with counts per million greater than zero, which ranged from 13,351 in the Liver to 15,959 in the VAT. Biological replicates across all tissues and the relationship between the tissues were analyzed using hierarchical clustering with pairwise Pearson correlation. We observed high Pearson correlation (0.92 to 1) between samples from the same tissue but different mice, suggesting that inter-individual variation has little impact on the transcriptomic profiles and demonstrating high reproducibility of the data, both at room temperature (RT, red section) and at cold exposure (CE, blue section) (Figure 2B). Larger clusters, as for instance the one between SAT, BAT, and VAT, both at RT and CE, indicate that physiologically close tissues show high similarity in terms of their global gene expression profiles.

**Figure 2:**
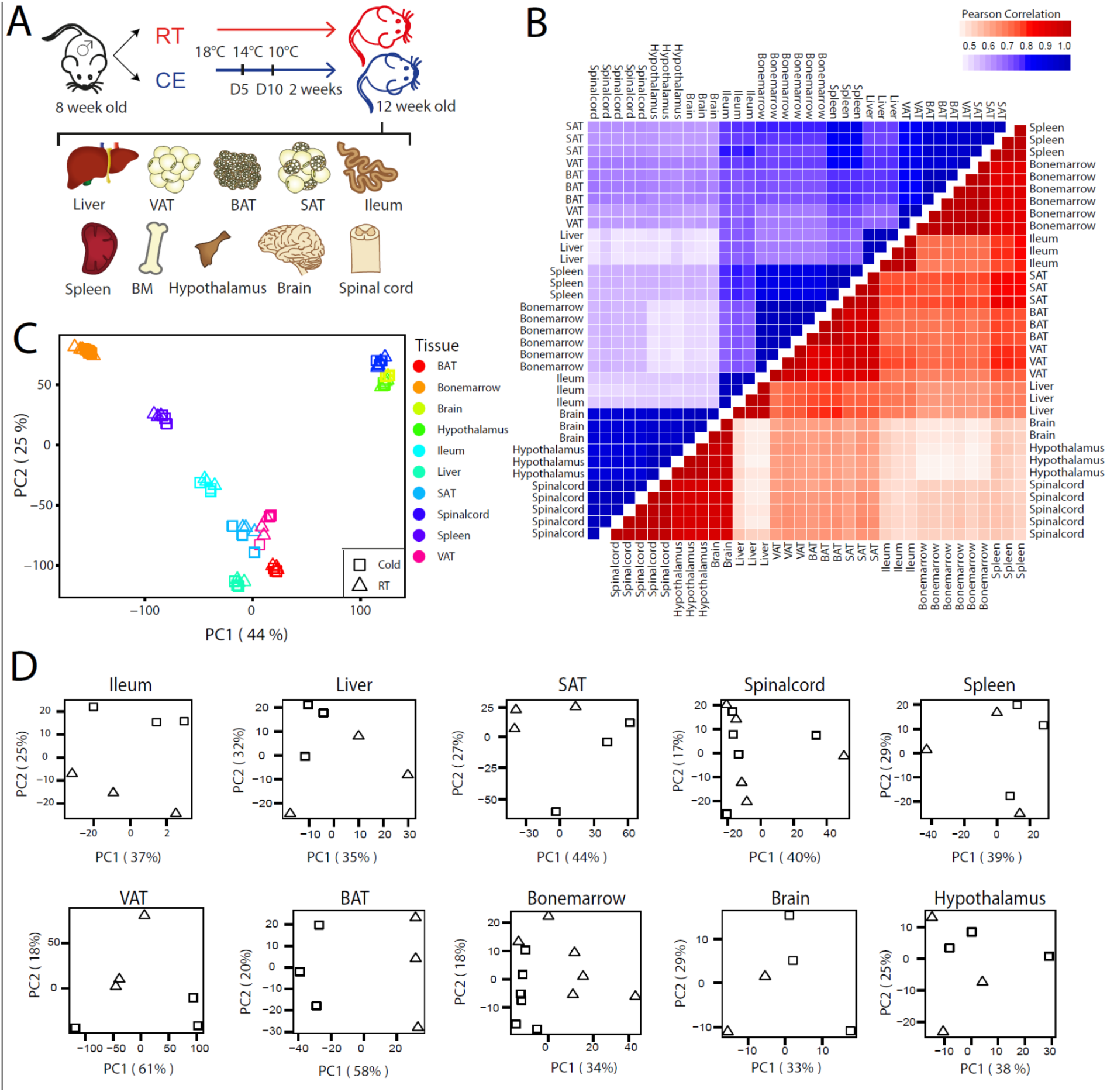
QC and clustering of the biological samples and tissues at cold and room temperature. (**A**) Experimental setup: 8 week old C57BL/6J mice were exposed to 10°C cold (CE) or room temperature (RT) for 2 weeks with initial 10 days of acclimatization and their tissues were harvested for RNA sequencing. Visceral adipose tissue (VAT), brown adipose tissue (BAT), subcutaneous adipose tissue (SAT), bone marrow (BM). (**B**) Correlation heatmap of 35 samples from the ten tissues of RT (red) and CE (blue) mice as in (A). Pearson’s correlation coefficient is computed for each pair of samples. (**C**) Principal component analysis (PCA) of the transcriptome across the ten tissues at RT and CE of mice as in (A). Physiologically close tissues are distinguished by global gene expression patterns. (**D**) PCA of samples for each tissue from mice as in (A).

To further investigate the clustering between the tissues, we performed a principal component analysis (PCA), using a combined dataset with 11403 genes that were identified across all ten tissues (RT and CE, Figure 2C). PCA revealed two strong clusters of physiologically related tissues (Figure 2C): The adipose tissue cluster with BAT, SAT, and VAT, and the CNS cluster with hypothalamus, brain and spinal cord samples. The bone marrow samples clustered relatively close to the spleen samples, which can be considered as an immune tissue cluster. The liver samples gathered close to the adipose tissues, while the ileum samples clustered between adipose tissue and spleen samples. Generally, the samples clustered rather by tissue and not by treatment (RT vs. CE).

To understand the propensity of samples from cold-exposed and RT-kept mice per given tissue, we performed PCA on each tissue separately (Figure 2D) and observed a different degree of clustering of RT and CE samples in the various tissues. Specifically, we detected a pronounced separation of RT and CE samples in BAT, opposed to a milder separation in hypothalamus samples, implying that the cold exposure has varying effects depending on the tissue.

### Deregulated genes and enriched GO biological terms upon cold exposure across the ten tissues

To further characterize the response to the cold exposure, we first assessed the number of deregulated genes and the magnitude of their deregulations across different tissues (Figure 3, Table S3), followed by GO gene set enrichment analysis of the identified differentially expressed genes. Collectively in all ten tissues, upon cold exposure we found 2471 genes as up-, and 3163 genes as down-regulated (p-value <0.05, |FC| > 1.5), while 679 genes were inversely regulated, i.e., upregulated in a given tissue(s) and downregulated in others. This inversely regulation may indicate that the regulation of certain genes occurs in a tissue specific manner. Physiologically close tissues, which clustered together (Figure 2C), shared similar numbers of deregulated genes. SAT underwent the most transcriptomic changes upon exposure to cold with 2867 significantly deregulated genes, followed by BAT and VAT, while brain and hypothalamus showed the least transcriptomic changes with 119 and 113 total deregulated genes, respectively (Figure 3). These results suggest that although adipose tissues are the most affected by cold exposure, all organs undergo changes that are reflected in their tissue-specific transcriptomic profiles.

**Figure 3:**
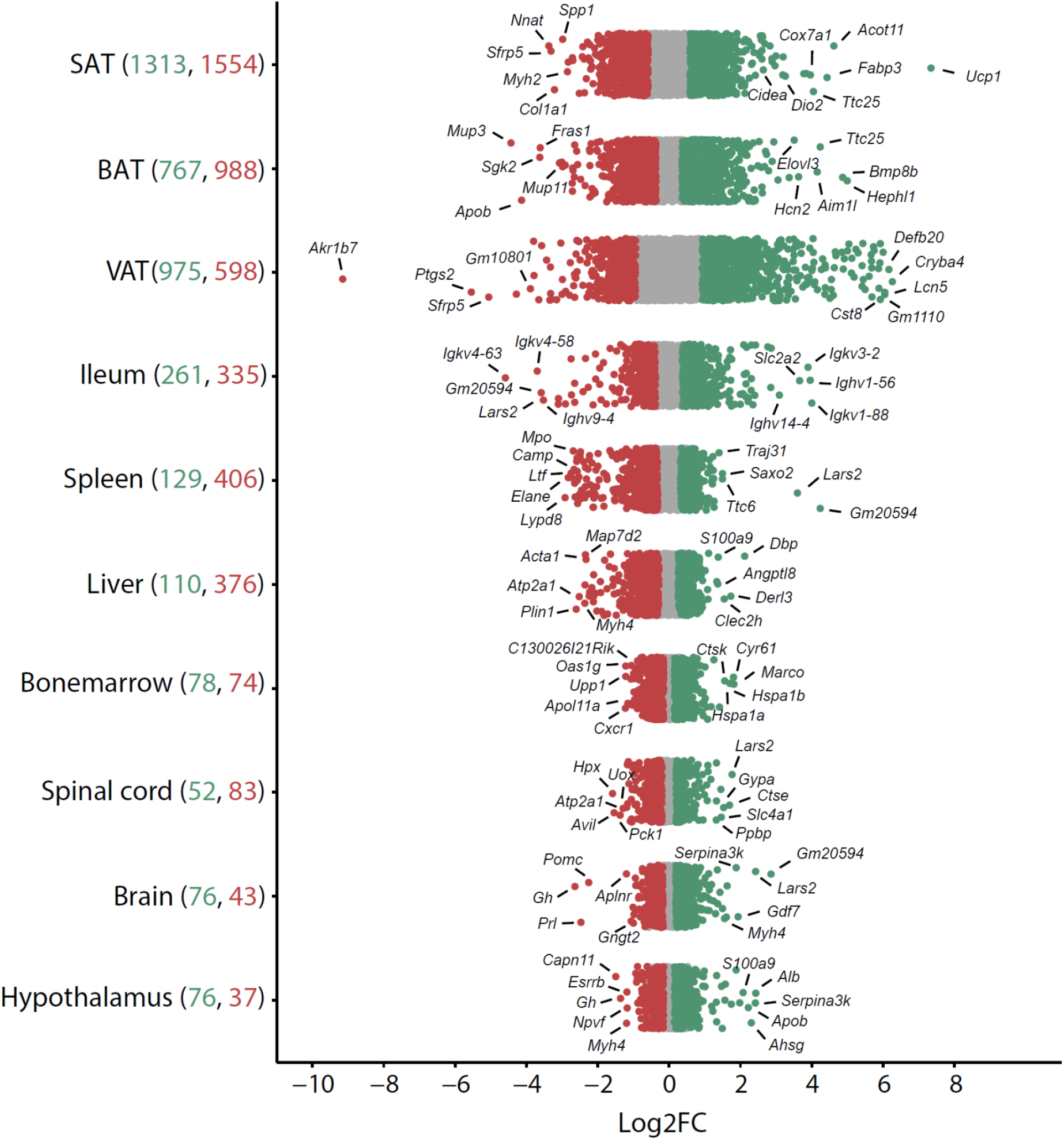
Distribution and degree of gene deregulation across ten tissues upon cold exposure. Log2FC of all genes across the ten tissues of mice as in Figure 2A (red and green colored dots indicate *P*<0.05, whereas grey dots indicate *P*>0.05). Numbers in parenthesis on the x-axis indicate the number of significantly up- (green) and down- (red) regulated genes. Gene name is shown for the five up- and five down-regulated genes with highest log2FC.

Moreover, the magnitude of gene deregulation (log2FC) showed an analogous ordering depending on the tissue. The adipose tissues had the biggest deregulation degree, including genes upregulated with log2FC of 7, e.g., Ucp1 in SAT and downregulated genes with log2FC of −9, e.g., *Akr1b7* in VAT, while spinal cord, brain and hypothalamus scored the mildest deregulations (Figure 3, Table S3).

We next performed GO gene set enrichment analysis on the deregulated genes for the ten tissues upon cold (|FC| > 1.5 and *P* -value < 0.05). Enriched gene sets with *P*-value < 0.01 were chosen for further analysis, collectively consisting of 434 downregulated and 259 upregulated gene sets (Table 1).

**Table 1:** GO-based gene set enrichment analysis across ten tissues upon cold exposure

| Significantly enriched GO terms (p-value <0.05) |  |  |  |
| --- | --- | --- | --- |
| Tissue | # of enriched GO terms | upregulated | downregulated |
| SAT | 261 | 83 | 178 |
| VAT | 115 | 42 | 73 |
| BAT | 96 | 43 | 53 |
| Spleen | 56 | 1 | 55 |
| Hypothalamus | 44 | 30 | 14 |
| Ileum | 37 | 25 | 12 |
| Liver | 30 | 3 | 27 |
| Bone marrow | 24 | 18 | 6 |
| Brain | 17 | 6 | 11 |
| Spinal cord | 13 | 8 | 5 |

SAT showed the highest degree of enrichment (261 GO terms), followed by VAT and BAT. The least changed tissues were spinal cord and brain with 17 and 13 deregulated gene sets, respectively (Table 1). Interestingly, this ordering was similar to the degree of gene deregulation (shown in Figure 3), with the exception of the hypothalamus, which ranked 5^th^ when the GO terms were used as criteria.

### Tissue-specific and tissue-shared gene and GO term signatures in response to cold exposure

To gain initial insights into the tissue-shared and tissue-specific enriched GO terms, the top 63 up- and 86 down-regulated terms (out of the 545 unique enriched GO terms, full list is shown in Table S4) are shown in Figure 4. The selection is made based on the *P* value in response to cold including both tissue-shared and tissue-specific pathways, and it concludes with the last pathway that shows tissue-shared response. Since this visual representation indicates that some cold responsive gene sets are shared between tissues, we next sought to unravel the genes and biological GO terms that exhibit tissues-specific, or common regulation patterns upon cold exposure across the ten tissues. We classified the deregulated genes and gene sets into three categories according to the number of tissues wherein they are differentially expressed upon cold exposure.

**Figure 4:**
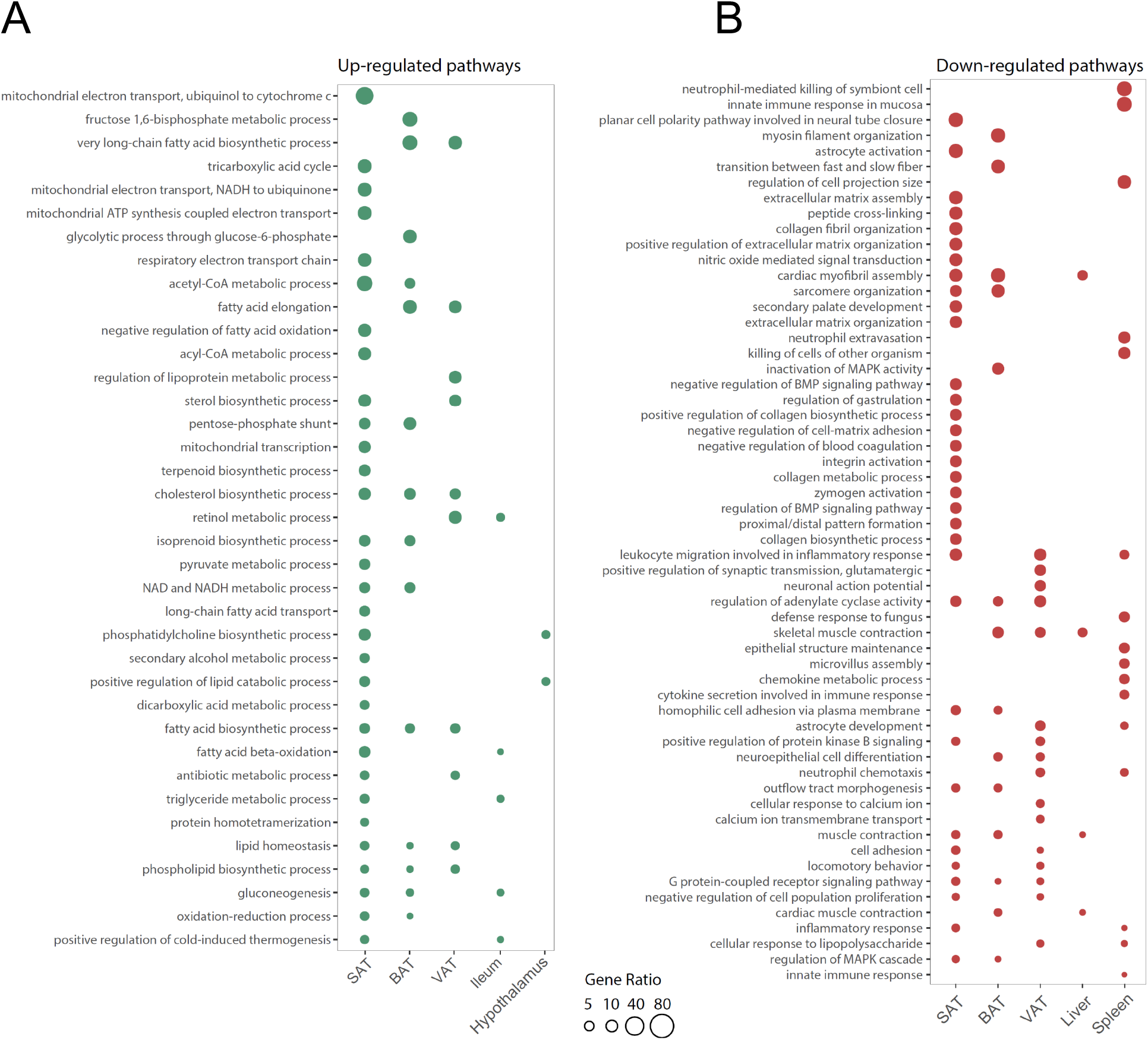
Visualization of highly enriched gene sets indicates common enrichment within groups of tissues upon cold exposure. **(A)** The top ranked 63 up-regulated gene sets out of 259, and **(B)**the top ranked 86 down-regulated gene sets out of 4 are visualized. The p-value and the gene ratio (the ratio of deregulated genes to annotated genes in each gene set) were used as the ranking criteria.

Altogether 2597 genes were upregulated in only one tissue (69% of the total upregulated genes) and were defined as “tissue-specific up-regulation profiles” and 2696 genes were downregulated in a single tissue (50% of the total downregulated genes) and were identified as “tissue-specific down-regulation profiles” (Table 2). We observed widespread tissue-specific cold response in the adipose tissues where SAT ranked highest with 921 up- and 1003 and down-regulated genes. We further compared the tissue-specific deregulated genes with the total number of deregulated genes for each tissue, which is indicated as a ratio in Table 2. The highest percentage of tissue specificity in terms of deregulated genes was seen in VAT with 77% tissue-specific upregulated genes, followed by spleen with 74% tissue-specific downregulated genes. Considering the total number of deregulated genes (sum of up and down regulated gens), SAT, VAT and spleen showed highest gene deregulation specificity with an average of 67% tissue-specific deregulation profiles (Table S3). Collectively, there was a higher degree of tissue-specificity in upregulated versus downregulated profiles (69% compared to 50%).

**Table 2:** Tissue-specific deregulated profiles. Number of up- and down-regulated genes per tissue shown as tissue-specific up-regulation profile (left) and tissue-specific down-regulation profile (right). The percentage of tissue-specific changes versus the total number of deregulated genes (as in Figure 3) is displayed in parenthesis.

|  | Tissue-specific upregulation profiles | Tissue-specific downregulation profiles |
| --- | --- | --- |
| **Tissue** | **# of genes (% of tissue-specific)** | **# of genes (% of tissue-specific)** |
| SAT | 921 (70%) | 1003 (65%) |
| VAT | 746 (77%) | 260 (43%) |
| BAT | 409 (53%) | 578 (59%) |
| Ileum | 172 (66%) | 218 (65%) |
| Spleen | 93 (72%) | 302 (74%) |
| Liver | 79 (72%) | 214 (57%) |
| Bone marrow | 53 (68%) | 47 (64%) |
| Hypothalamus | 50 (66%) | 25 (68%) |
| Brain | 47 (62%) | 27 (63%) |
| Spinal cord | 27 (52%) | 22 (27%) |
| **Total** | **2597 (69%)** | **2696 (50%)** |

We next focused on the genes with shared deregulation profiles within several tissues. Interestingly, there was no overall common gene regulation set for all ten tissues. The maximum tissue-shared gene signature belonged to *Atp2a1*, which was decreased in 7 tissues (All adipose tissues, liver, bone marrow, hypothalamus, and spinal cord). The next top tissue-shared gene signatures were 6 genes that were commonly downregulated (*Myh4, Nnat, Nr1d1, Tnni2, Tnnt3*) and upregulated (*Thrsp*) in 5 tissues. The rest of the tissue-shared genes were shared within groups of four, three, and two tissues. To illustrate the tissue-shared genes and their distribution within the ten tissues, we performed a network analysis (Figure 5 A & B), where the size of each node is correlated with the number of tissue-specific cold-deregulated genes of each tissue, e.g., the SAT node is the biggest as it represents 921 upregulated (Figure 5A, Table 2) and 1003 downregulated (Figure 5B, Table 2) genes.

**Figure 5:**
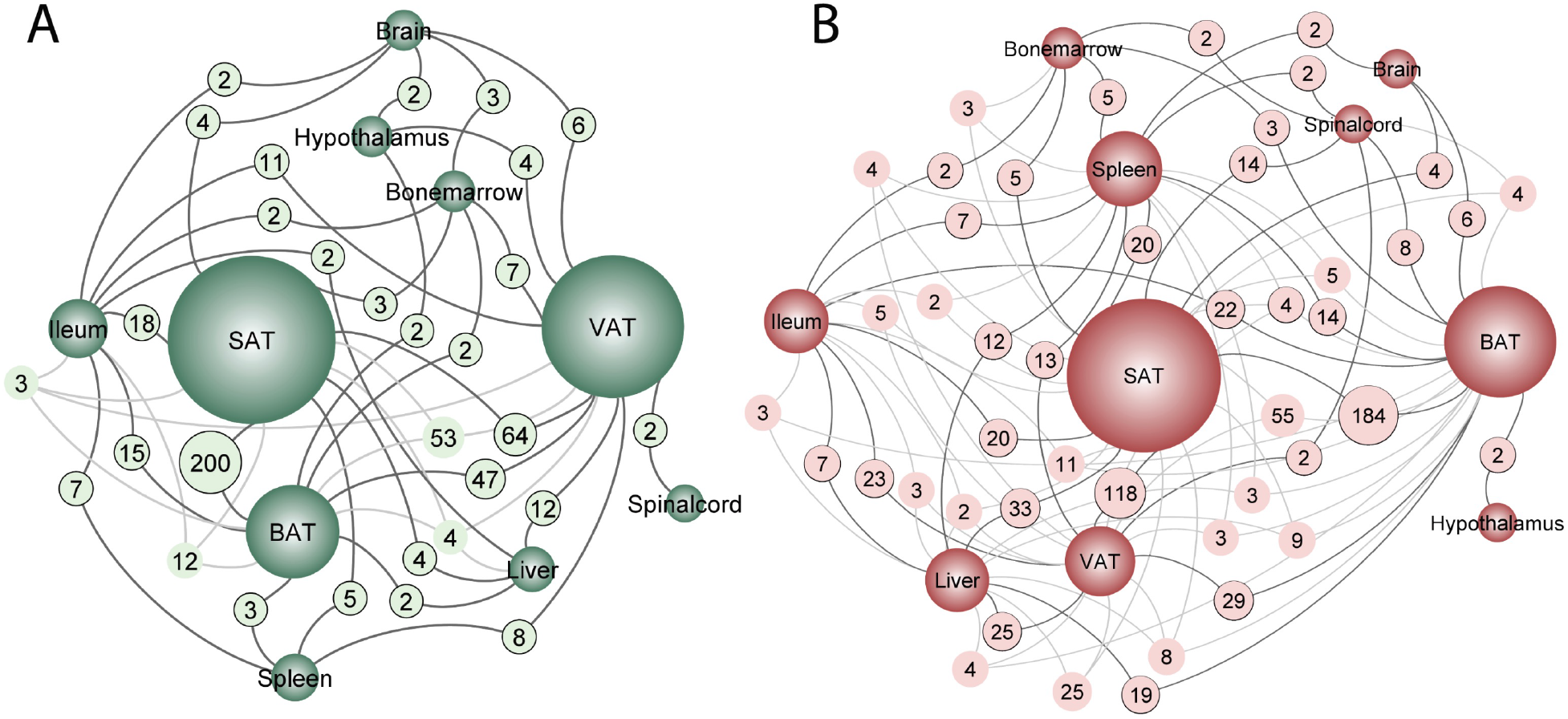
Tissue-shared gene deregulations upon cold exposure. Network plot visualizing the relationship of tissue-shared (numbered light green and light red circles) and tissue-specific (dark green and dark red circles) genes in the ten tissues of mice, as in Figure 1A. Upregulated genes upon cold exposure are shown in the green network (**A**), and downregulated genes are shown in the red network (**B**). Tissue-shared genes with a minimum of two shared genes are displayed, and the full dataset is provided in Table S3. Tissue-specific genes are connected to tissue-shared genes (light green and light red) with dark grey lines if they indicate a pairwise connection and in light grey, if they are shared in more than two tissues. The size of each node indicates the number of both shared and specific deregulated genes.

The highest degree of tissue-shared upregulated genes was found within the adipose tissues, where 200 genes were similarly upregulated in SAT and BAT, followed by 64 genes in VAT and SAT and 47 genes in VAT and BAT (Figure 5A). Likewise, SAT, VAT and BAT shared 53 genes, which are upregulated in the three tissues. Apart from the adipose tissues, the top-shared pairwise connections were related to ileum and liver. The analysis also revealed that spinal cord shares the least number of upregulated genes with other tissues, having only two genes shared with VAT.

Similarly, the top-shared pairwise connections for downregulated genes were found in adipose tissues where SAT and BAT shared 184 downregulated genes, SAT and VAT 118 genes, followed by liver and SAT with 33 tissue-shared downregulated genes. The hypothalamus displayed lowest number of shared downregulated genes, having only two in common with BAT.

Next, we studied the tissue-specific and the tissue-shared GO terms enrichment. Out of the 545 regulated GO terms, 445 (81%) were identified as tissue-specific regulated pathways (Table S4). This suggests that the transcriptional cold response at the level of functional gene sets is to a great degree tissue-specific. To unravel which cold responses are shared between tissues, we identified commonly regulated gene sets. Of note, the maximum number of tissues which shared a given regulated gene set is three. A closer inspection of the 63 commonly deregulated gene sets revealed that 38 were commonly downregulated between three (6 gene sets) and two (32 gene sets) tissues, while 25 gene sets were commonly upregulated between three (5 gene sets) and two (20 gene sets) tissues (Figure 4, Table 3, and the full list in Table S3). We observed a clear overrepresentation of the adipose cluster in the shared deregulated gene sets, which suggests a common response in SAT, BAT, VAT that extends to ileum, liver and spleen for certain GO terms. This indicates that the closely related tissues share higher numbers of cold responsive gene sets.

**Table 3:** Shared tissue GO gene set enrichments within three tissues upon cold exposure

| GO gene set enrichment | Tissues |
| --- | --- |
| Shared tissue gene set enrichments within upregulated genes (p-value <0.05) |  |
| fatty acid biosynthetic process | SAT,BAT,VAT |
| phospholipid biosynthetic process | SAT,BAT,VAT |
| cholesterol biosynthetic process | SAT,BAT,VAT |
| lipid homeostasis | SAT,BAT,VAT |
| gluconeogenesis | SAT,BAT,ileum |
| Shared tissue gene set enrichments within downregulated genes (p-value <0.05) |  |
| G protein-coupled receptor signaling pathway | SAT,BAT,VAT |
| muscle contraction | SAT,BAT,liver |
| leukocyte migration involved in inflammatory response | SAT,spleen,VAT |
| cardiac myofibril assembly | SAT,BAT,liver |
| regulation of adenylate cyclase activity | SAT,BAT,VAT |
| skeletal muscle contraction | BAT,liver,VAT |

### Shared-tissue regulated GO terms involve tissue-specific and tissue-shared genes

We next investigated whether the tissue-shared GO terms regulation is derived from the shared or the tissue-specific deregulated genes. We particularly focused on SAT, VAT and BAT. The Venn diagram of the first three most commonly up- and down-regulated gene sets showed that they are largely derived by genes that are deregulated in a tissue-specific manner (Figure 6A). This tissue specificity is more pronounced in the downregulated gene sets (Figure 6B). For example, in fatty acid biosynthesis process, only 4 genes are shared between the three tissues and the upregulation of the pathway mostly emerges from the tissue-specific genes in SAT (13 genes), VAT (9 genes) and BAT (7 genes). This suggests that although different genes are up-regulated in the three tissues, they might have a similar function that leads to a common functional response. Interestingly, the degree of tissue-specific gene upregulation is on average higher for VAT followed by BAT and SAT.

**Figure 6:**
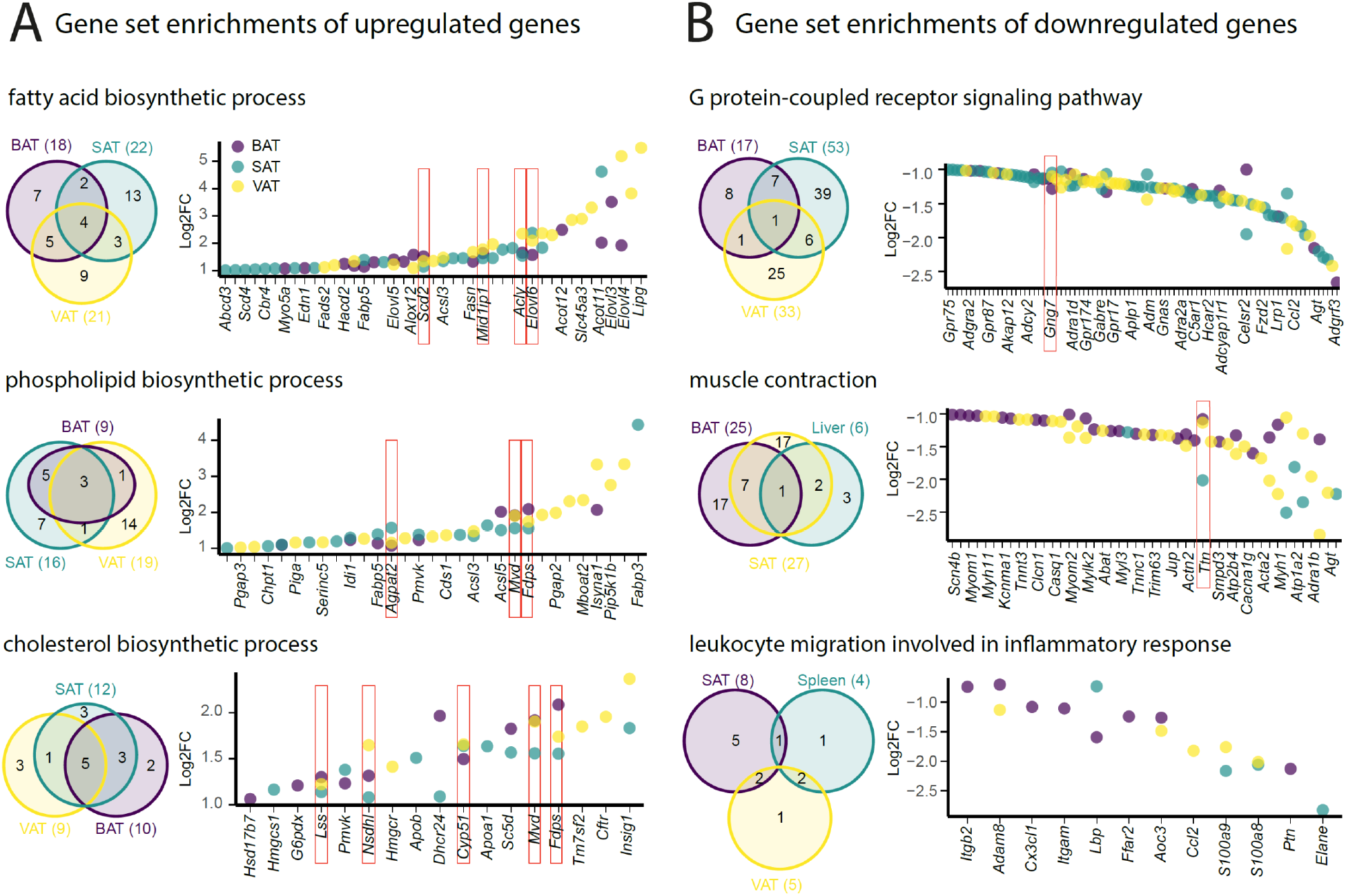
Shared tissue GO term enrichments and implicated genes. **(A-B)** Venn diagram of shared adipose tissue GO gene set enrichments of up- (A) and down- (B) (left), and dot plot of genes involved in these gene sets (right) across the adipose tissues from mice as in Figure 2A. Not all the genes are labeled on the dot plots.

### Tissue-specific response to cold is not orchestrated by the gene expression patterns

To investigate if the strong tissue specificity in response to cold exposure is directed by the tissue specificity in the overall gene expression patterns, we first accounted for the genes with minimum 5 raw counts as a filtering threshold, and observed that 19,747 genes are expressed at least in one tissue, wherein 10977 genes (55% of expressed genes) were expressed in all ten tissues (Figure 7A). From these, 3407 genes (36%) were deregulated in at least one tissue (Figure 7B). Looking at these deregulated genes, irrespective of the consistency in the direction of change, we found that 2428 genes (over 71%) were altered in only one tissue, 731 (over 21%) in two, and 190 (over 5%) commonly deregulated in three tissues. The rest 1.7% of deregulated genes were commonly changed in four or more tissues (Figure 7C). These data demonstrate high tissue-specificity in the transcriptional response to cold also among the commonly expressed genes. Moreover, 1623 genes (8 % of expressed genes) were expressed in only one tissue (Figure 7A). Spinal cord with 554 genes showed the most tissue-specificity in term of expressed genes, followed by VAT (339 genes), bone marrow (202 genes), ileum (140 genes), spleen (138 genes), liver (96 genes), hypothalamus (79 genes), brain (52 genes), BAT (13 gens) and SAT (10 genes). Interestingly, only 66 genes of the 1623 tissue-specific expressed genes were significantly deregulated upon cold exposure (Figure 7B). Together, this indicates that the high degree of tissue-specificity in the transcriptional responses during cold exposure is not orchestrated by the global expression pattern exhibited by the various organs.

**Figure 7:**
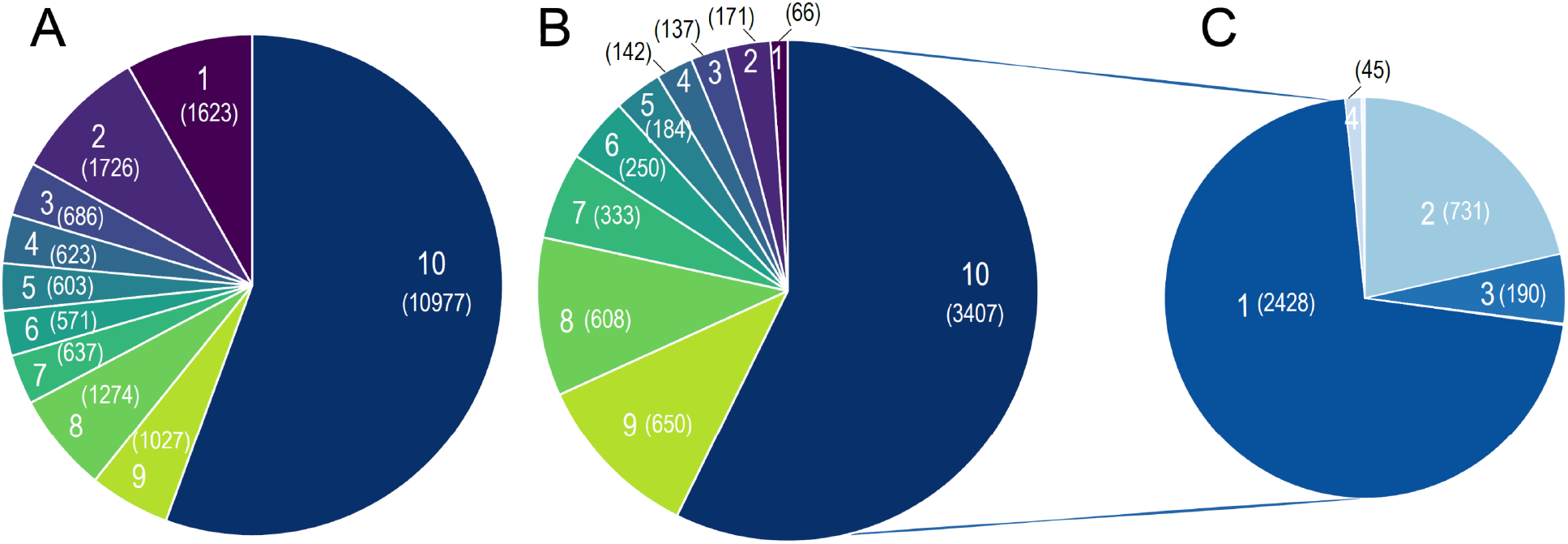
Global tissue-shared and tissue-specific gene expression and regulation patterns across ten tissues during cold. (**A**) Pie chart depicting distribution of the 19,747 genes on the number of tissues. Number of tissues where is indicated on each slice of the Pie chart, and the number of commonly expressed genes for the respective number of tissues is shown in parenthesis, e.g., 10,977 genes are expressed in all ten tissues and 1,623 genes are expressed in only one tissue (**B**) Distribution of 5,948 deregulated genes during cold based on the number of tissues. Number of tissues is indicated on each slice of the Pie chart and the number of genes is shown in parenthesis. Number of tissues and colors are kept in the same order as in (A), e.g., out of the 1623 genes that are expressed only in one tissue, 66 genes are deregulated. (**C**) Distribution of 3,407 that are expressed in all ten tissues (as in B) that are deregulated in at least one tissue. The number shows number if tissues where the gene is deregulated, and in parenthesis are the number of deregulated genes per respective number of tissues. E.g. 2,428 genes are deregulated in only one tissue.

To further underpin these conclusions, and given the importance of the thermogenic and fatty acid metabolism biomarkers in the cold response, we specifically analyzed and compared the tissue specificity on both global expression and regulation levels of the 148 thermogenic and 292 fatty acid metabolism biomarkers (Table S1 & Table S2) across the ten tissues (Figure 8). All the 148 thermogenic markers are expressed (minimum threshold of 5 raw counts) at least in two tissues, i.e., there is no gene that is specifically expressed only in one tissue. Surprisingly, we found only 30 genes which are not expressed in all ten tissues, which is in agreement with the global expression analysis. Liver contributes the most to this list by not expressing 16 of these 30 genes (Figure 8A, Table S5). Although all the 148 thermogenic biomarkers are expressed across most of the ten tissues, only 97 genes exhibit significant deregulation (p-value <0.05, |FC| > 1.5) upon cold exposure (Figure 8C), wherein 71 belong to adipose tissues. SAT has the highest number deregulated thermogenic biomarkers, followed by BAT and VAT (65, 31 and 25 deregulated genes, respectively), while the brain showed no thermogenic biomarker deregulation (Figure 8C).

**Figure 8:**
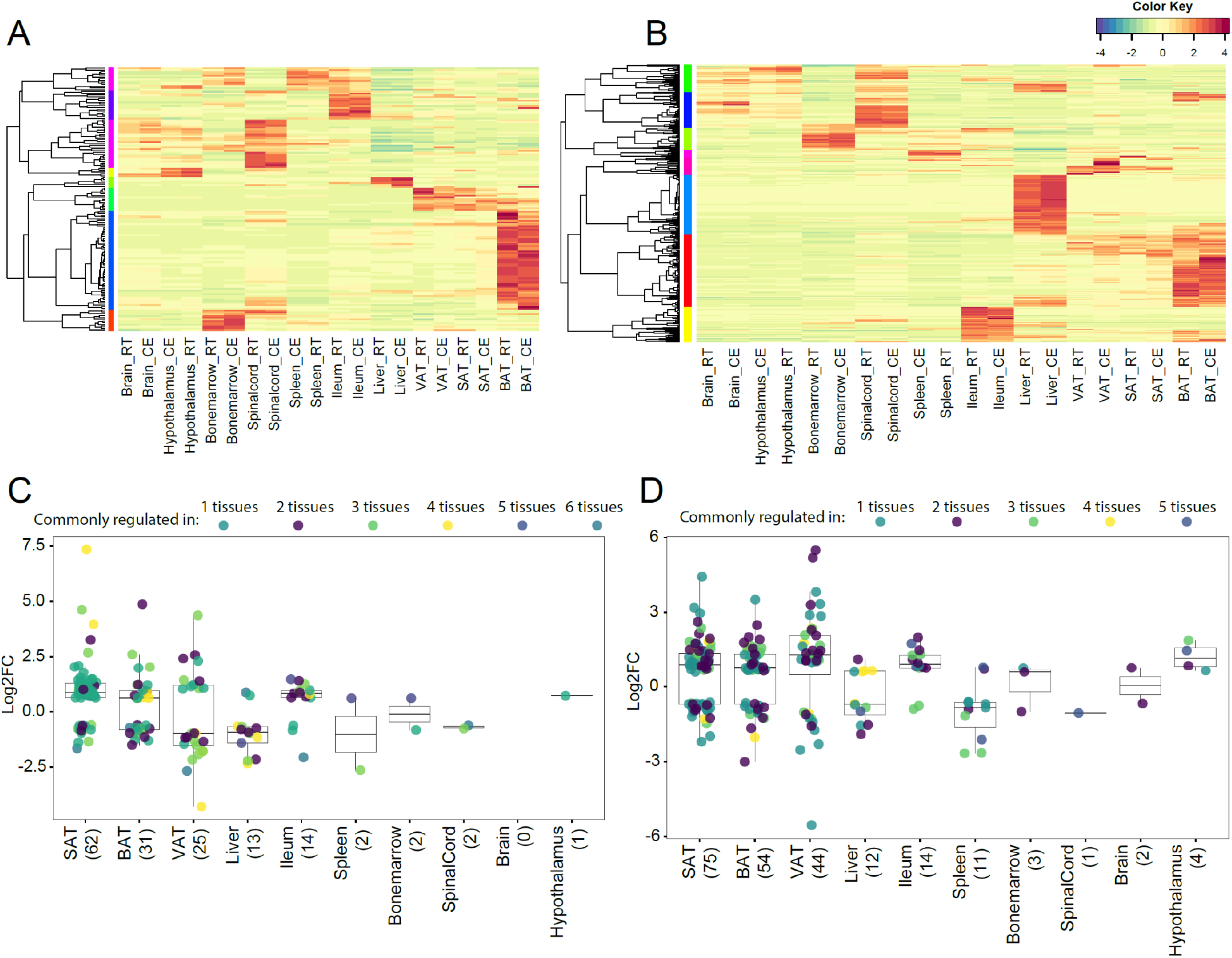
Expression and regulation of thermogenic and fatty acid metabolism biomarkers across ten tissues. (**A**) and (**B**) Heatmap showing an average of the thermogenic and fatty acid metabolism biomarker expression values of mice at RT or cold as in Figure 2A across the ten tissues. (**C**) and (**D**) Log2FC of the significantly (*P*<0,05) deregulated thermogenic and fatty acid metabolism biomarkers across the ten tissues. Each dot corresponds to the log2FC of a gene and colors specify the number of tissues where that gene is deregulated. The total number of deregulated genes for each tissue is shown in parenthesis.

Out of the 292 fatty acid metabolism biomarkers, 278 genes are expressed (minimum threshold of 5 raw counts) in at least one tissue. Similarly, most of the expressed genes (75 %) were expressed in the ten tissues and we did not observe a strong tissue specificity on the level of gene expression in fatty acid biomarkers (Figure 8B). 137 genes out of the 278 expressed genes were deregulated in at least one tissue. We identified 83 tissue-specific gene deregulations, where within SAT we found 37 tissue-specific deregulated genes ranking it on the top, followed by BAT with 23 genes and VAT with 16 genes. The spinal cord exhibits no tissue-specific deregulation of fatty acid metabolism biomarkers (Figure 8D, Table S6). The strong tissue-specificity pattern of deregulated genes (Figure 5) was further observed in the thermogenic (Figure 8C) and the lipid metabolism (Figure 8D) biomarkers. Collectively, these results suggest that tissue specificity is dictated in the response to cold (gene regulation level) and not on the gene expression level.

### Tissue-specificity of genes from the same family and with similar functions

By grouping the 19,747 expressed genes of our mouse gene catalog solely based on their gene nomenclature and without considering their specific function, we identified 2827 belonging to a “group of genes”. Members of these groups range from 1084 genes in the biggest group - *Olfr* (Olfactory receptors), to 2 genes in the smallest group (1273 of these groups have only two members) (Table S7). We found several thermogenic and fatty acid metabolism biomarkers such as *Abhd, Abhd, Acsm, Pex, Ppar, Hacd, Elovl, Apoa*, etc. as protein families, where we performed comparative analysis of their gene expression and regulation across the ten tissues (Table S5, Table S6). Similar as before, we observed tissue-specificity on the response to cold, but not on the gene expression patterns. As an example, we describe the elongation of very long chain fatty acids (*Elovl*) protein family, which plays an important role in fatty acid metabolism in adipose tissues. *Elovl* family includes seven genes named *Elovl1* to *Elovl7*, and catalyzes the first and rate-limiting reaction of the very long-chain fatty acid elongation cycle in the fatty acid biosynthesis pathway (Jump, 2009). The seven members displayed differing gene expression levels (Figure 9A) and variable rates of deregulation upon cold exposure across the ten tissues (Figure 9B). *Elovl1* and *Elovl7* are present at all ten tissues with particularly high expression in Ileum, but without any significant change upon cold exposure. The liver expresses all genes except *Elovl7* and *Elovl4* at a very high level, however we found no significant change in the expression levels at cold. The highest average expression value belongs to *Elovl6* in BAT and the highest regulation was seen for *Elovl3* in SAT. Elovl5 and Elovl6 expression levels were relatively high across all tissues and showed a pronounced upregulation by cold in the three adipose tissues (Figure 9B). Moreover, we observed a VAT-specific increase in *Elovl2* expression upon cold. *Elovl3* on the other hand was upregulated in BAT and SAT, whereas *Elovl4* expression was enhanced in BAT and VAT from the cold-exposed mice compared to their RT-kept controls and the magnitude of upregulation was much higher in VAT (Figure 9B). Consistent with our previous analysis, although different members of *Elovl* gene family were expressed in most of the tissues, their response to cold was tissue specific.

**Figure 9:**
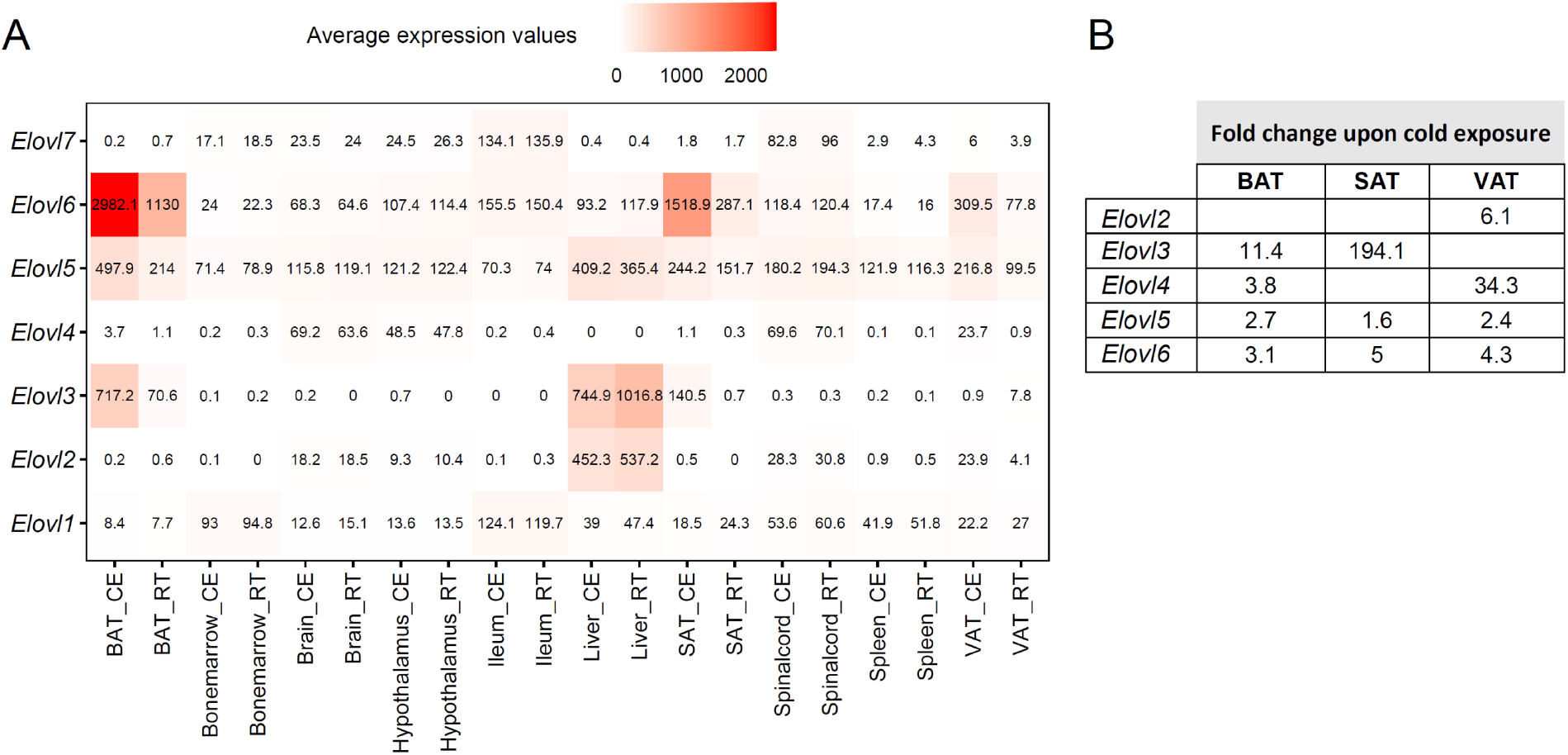
Average expression and regulation of Elovl gene family across ten tissues at RT and cold. (**A**) Heatmap showing average of the normalized gene expression values of mice at RT or cold as in Figure 2A across ten tissues. (**B**) Log2FC of the significantly upregulated genes (*P*<0,05) of the *Elovl* family from mice as in (A). Significant changes of the *Elovl* family were detected only in in the adipose tissues.

## DISCUSSION

Cold exposure is an extensively studied intervention to promote BAT activity and SAT browning. Despite the potential therapeutic relevance of this environmental trigger in increasing the energy expenditure, the impact of cold on the transcriptomic landscape of other tissues and their physiological response has received little attention so far. On the other hand, different temperatures have been interchangeably used in cold exposure studies. In this study, we first gathered all publicly available RNASeq profiles of C57BL/6J mouse BATs obtained under different cold exposures ranging from 4 °C to 8 °C. This comparative analysis revealed that biomarker response to our milder (10°C) cold setup was effectively in the 50^th^ percentile range with the exception of a few genes e.g., *Dio2*. Interestingly, our results indicate that the variance in magnitude of the transcriptional upregulation across the different studies can be explained by the duration of the cold exposure rather than the extent of the temperature decrease *per se*, as evidenced by the expression of the thermogenic and fatty acid metabolism biomarkers.

When analyzing the comparative RNAseq data across the ten tissues, we observed that as expected, similar tissues effectively clustered together in terms of their overall transcriptomic profile, regardless of the imposed condition, i.e., cold exposure or RT. The extent of the sample separation following cold was different for different tissues, suggesting that cold exposure causes variable and tissue-specific responses. These conclusions were further supported by the number and the magnitude of deregulated genes and the enriched gene sets. Upregulated GO terms, which were mostly related to thermogenesis, oxidative phosphorylation, fatty acid biosynthesis, elongation and catabolic processes, and mitochondrial ATP synthesis scored the highest in the adipose tissues. This was followed by the hypothalamus and the ileum where some of the upregulated pathways related to lipid metabolism went in the same direction as the adipose tissues, however to a lower extent. Strikingly, the tissue specificity in the response to cold was not dictated by the gene expression differences between the various tissues, since we found the same levels of alterations even within the commonly expressed genes (55%).

The multi tissue data in this study opens the possibilities for integrative analyses of the adaptation to lower environmental temperatures, and it allows investigating how organs compensate for the increased activity of the above-mentioned biological process. As expected, in the adipose tissues, which showed the highest number of upregulated gene sets particularly related to thermogenesis, we identified a considerable number of downregulated gene sets. This may suggest that on a tissue level the energy expenditure is balanced by a tendency for lowering a range of biological processes to maximize the necessary biological response to a given trigger.

On the other hand, such balance between up- and down-regulated biological processes was not seen in the spleen, where we observed primarily downregulated gene sets. Several biological processes such as innate immune response in mucosa, neutrophil-mediated killing of symbiont cell, inflammatory response, and many other responsive systems, e.g., response to bacteria, were all highly downregulated in spleen. The predominant down-regulations in the spleen could be explained by its mainly immunologic functions, a biological process that might be blunted during cold. Intriguingly, our results further indicated that some pathways are inversely regulated across tissues. For example, the triglyceride catabolic process was upregulated in SAT, brain, and hypothalamus, but downregulated in spleen. In part, this may indicate an overall redistribution of the metabolic energy towards the tissues necessary for acute response to an environmental trigger such as cold, on account of the other maintenance biological programs.

Examining tissue-specific and tissue-shared genes and gene set signatures revealed that tissue-specificity both at level of genes and gene sets dominate over tissue-shared patterns. Interestingly, even within the adipose tissues, we observed preferential tissue-specific response over shared characteristics. Interestingly, the shared-tissue gene sets, e.g., upregulation of fatty acid biosynthetic process, do not necessarily require upregulation of the same genes. This highlights the tissue specificity also on a gene level, where regulation of different genes contributes to emergence of the same biological process. Similarly, analysis of the *Elovl* gene family in context of the different tissues revealed that different members of a given gene family could evolve and be regulated in a tissue-specific manner. Such inter-tissue specificity of genes from the same family can guide discovery of tissue-specific regulators upon cold adaptation and contribute to a better understanding of the underlying molecular basis of cold adaptation of each tissue. In summary, our work shows that 10°C cold exposure causes a representative transcriptomics response in BAT, and highlights the local alternations in the transcriptomic profile across ten tissues, which include neural, immune, and metabolic responses. We believe that this tissue-specific cold-induced expression atlas will be a useful resource for studding the physiological alterations in response to lower environmental temperatures in an integrative manner.

## METHODS

### Mice

Male 8 weeks old C57BL/6J mice were obtained from Janvier (France). Mice were housed in a specific pathogen free (SPF) facility in 12 h day/night cycles with free access to irradiated standard chow diet and water from autoclaved bottles. All mice were housed 2 per cage without bedding and nesting material to ensure controlled temperature conditions. Cold exposure was performed in a light- and humidity-controlled climatic chamber (TSE, Germany) under SPF conditions, at 10 °C for 2 weeks with an initial acclimatization period of 5 days at 18°C and 5 days at 14°C. All animal experiments were performed at the Universities of Geneva with authorization by the responsible Geneva cantonal, and Swiss federal authorities in accordance with the Swiss law for animal protection.

### Experimental model and subject details

Our study design is summarized in table S8. Transcriptome dataset were generated from ten different tissues (Brown adipose tissue (BAT), bone marrow, brain, hypothalamus, ileum, liver, inguinal subcutaneous adipose tissue (SAT), spinal cord, spleen, and epididymal visceral adipose tissue (VAT)) of C57BL/6J mice subjected to cold, or kept at RT. The summary of mice and tissue sampling is provided in Table S8.

### RNA extraction for RNA sequencing

After collection, all tissues except bone marrow were snap frozen in liquid nitrogen and stored at −80°C until used. Frozen tissues were mechanically homogenized with 1 stainless steel bead (5 mm) in 1 ml Trizol (Thermo Fisher Scientific) by shaking for 50 s at 30 Hz (TissueLyser, Qiagen). 200 μL chloroform was added to homogenize Trizol samples, followed by 15 s shaking and centrifugation (15 min, 12000 RCF, 4°C). The chloroform phase was collected, shaken for 15 s with 500 μL isopropanol and centrifuged (50 min, 12000 RCF, 4°C). The pellet was washed with 70% ethanol twice (10 min, 8000 RCF, 4°C) and dissolved in 50 μL PCR-grade water. Bone marrows were flushed immediately after collection from mouse, cells were spun and loaded onto shredder columns (Qiagen). Shredded bone marrow cells were frozen (−80°C) in RLT buffer (1% beta-mercaptoethanol) until RNA extraction using RNAeasy mini kit (Qiagen). RNA integrity number (RIN) was determined in all samples (Bioanalyzer 2100, Agilent) before sequencing.

### RNAseq sequencing

The mRNA sequencing was done at the iGE3 Genomics Platform at the CMU of the University of Geneva for the bone marrow and spinal cord, while for all other tissues at the University Medical Center Göttingen (UMG), Institute of Human Genetics, NGS-Integrative Genomics Core Unit (NIG). Libraries for sequencing of bone marrow and spinal cord were prepared with the TruSeq stranded mRNA kit and sequenced with read length SR50 (Illumina HiSeq 4000). For all other tissues RNA-seq libraries were prepared using the NEBNext Ultra RNA Library Prep Kit for Illumina, and were pooled and sequenced on an Illumina HiSeq 4000 sequencer generating 50 base pair single-end reads as in (Schattling et al., 2019, Di Liberto et al. 2018) with 30 Mio reads/sample. The sequencing quality control was done with FastQC v.0.11.5 (http://www.bioinformatics.babraham.ac.uk/projects/fastqc/). The reads were mapped with STAR aligner v.2.6.0c (Dobin et al., 2013) to the UCSC Mus musculus mm10 reference. The transcriptome metrics were evaluated with the Picard tools v.1.141 (http://picard.sourceforge.net/). The table of counts with the number of reads mapping to each gene feature of the Mus musculus mm10 reference was prepared with HTSeq v0.9.1 (htseq-count, http://www-huber.embl.de/users/anders/HTSeq/).

### Data analysis

Raw counts were processed and analyzed by R/Bioconductor package EdgeR v. 3.4.2 (McCarthy et al., 2012), for normalization differential expression analysis. The counts were normalized according to the library size and filtered. Only genes having log count per million reads (cpm)>0 were kept for the further analysis. After normalization of the counts, transcript abundances were compared in pairwise conditions in a modified Fischer exact test (as implemented in edgeR). Two-tailed unpaired Student’s t-test was used for pair-wise comparisons, and p <0.05 was considered statistically significant, unless otherwise specified. Genes were considered significantly changed if they passed a fold change (FC) cutoff of |FC| > 1.5 and a p-value≤0.05, and were further subjected to gene ontology analysis, using R/Bioconductor package topGO (https://bioconductor.org/packages/release/bioc/vignettes/topGO/inst/doc/topGO.pdf), together with Rgraphviz, Pearson correlation similarity analysis, and heatmap visualization. Principal Components Analysis (PCA) and volcano and dot plots were generated in R, and the scripts used for the analysis and generating figures are available at github.com/Nhadadi/Mouse_AllTissue_Transcriptomics.

### Network analysis

Network analysis of all genes in the tissue-enriched and group-enriched categories was done using Cytoscape 3.8 (Shannon et al., 2003). The resulting network includes only group enriched nodes with at least two expressed genes.

## SUPPLEMENTARY TABLE LEGENDS

**Table S1:** Log2FC of 148 potential thermogenic biomarkers combined from literature and GO database across 7 studies. Only significantly deregulated genes (*P*<0.05) are shown.

**Table S2:** Log2FC of 292 potential fatty acid metabolism biomarkers from GO database across 7 studies. Only significantly deregulated genes (*P*<0.05) are shown.

**Table S3:** Log2FC of significantly (*P*<0.05) deregulated genes across the ten tissues.

**Table S4:** Enriched GO terms across the ten tissues.

**Table S5:** Average of the normalized gene expression values of thermogenic and fatty acid metabolism biomarkers of mice at RT or cold across the ten tissues.

**Table S6:** Deregulation of thermogenic and fatty acid metabolism biomarkers across the ten tissues. Only significantly deregulated genes (*P*<0.05) are shown.

**Table S7:** Gene family members.

**Table S8:** Summary of the study design.

## DATA AVAILABILITY

The raw counts from the RNA-seq data, and the code for the bioinformatics pipeline developed for this study have been made freely available at (github.com/Nhadadi/Mouse_AllTissue_Transcriptomics). Prior to the publication, the RNAseq data will be also deposited to GEO and given accession number.

## ACKNOWLEDGEMENTS

We thank the iGE3 Genomics Platform at the Medical faculty of the University of Geneva, and the NGS-Integrative Genomics Core Unit (NIG) at the Institute of Human Genetics, University Medical Center Göttingen (UMG) for mRNA sequencing. This work was supported by the CONFIRM grant of the Hôpitaux Universitaires de Genève (HUG) Foundation (no. RC2-09) and the Swiss Multiple Sclerosis Society Grant to DM and MT, the European Research Council (ERC) under the European Union’s Horizon 2020 research and innovation program (ERC Consolidator grant agreement no. 815962, Healthybiota) to MT; and Swiss National Science Foundation (SNSF) grant to DM (310030_173010).

## AUTHOR CONTRIBUTIONS

N.H. set up the bioinformatics pipeline for the RNAseq and downstream data analysis. M.S. and K.S. set up and performed the mouse experiment and RNA isolations. G.S. performed the sequencing as specified in Methods. N.H. and M.S. analyzed and interpreted the results, and drafted figures. D.M. and M.T. initiated, designed and supervised the project. N.H., M.S and M.T interpreted the data wrote the manuscript. All authors read and approved the final manuscript.

